# An intriguing characteristic of enhancer-promoter interactions

**DOI:** 10.1101/2020.05.24.112458

**Authors:** Amlan Talukder, Haiyan Hu, Xiaoman Li

**Author notes:** To whom correspondence should be addressed. Correspondence may also be addressed to Xiaoman Li.

## Abstract

It is still challenging to predict interacting enhancer-promoter pairs (IEPs), partially because of our limited understanding of their characteristics. To understand IEPs better, here we studied the IEPs in nine cell lines and nine primary cell types. We observed that one enhancer is likely to interact with either none or all of the target genes of another enhancer. This observation implies that enhancers form clusters, and every enhancer in the same cluster synchronously interact with almost every member of a set of genes and only this set of genes. We perceived that an enhancer can be up to two mega base pairs away from other enhancers in the same cluster. We also noticed that although a fraction of these clusters of enhancers do overlap with super-enhancers, the majority of the enhancer clusters are different from the known super-enhancers. Our study showed a new characteristic of IEPs, which may shed new light on distal gene regulation and the identification of IEPs.

## INTRODUCTION

Enhancers are short genomic regions that can boost the transcription of their target genes under specific experimental conditions (1,2). They directly interact with the promoters of their target genes via chromatin looping to control the temporal and spatial expression of target genes (3-6). Enhancers can be several dozens to a couple of thousand base pairs (bps) long and can be located in the distal upstream or downstream of their target genes (1). Although the longest experimentally validated distance between enhancers and their targets is about one mega bps (Mbps), it is not rare that enhancers are more than two Mbps away from their target genes (5,6). Because of such a long distance, it is still challenging to identify interacting enhancer-promoter pairs (IEPs). In this study, an IEP refers to an enhancer-promoter pair that physically interacts, although such an interaction may or may not have any functional effect observed yet.

Many methods are available to identify enhancers. Early experimental studies identify enhancers by “enhancer trap”, which has established our rudimentary understanding of enhancers in spite of its low-throughput and time-consuming nature (7,8). Early computational methods predict enhancers through comparative genomics, which are cost-effective but may produce many false positives (9,10,46,49). With the next-generation sequencing (NGS) technologies, enhancers are identified through a variety of experimental methods such as chromatin immunoprecipitation followed by massive parallel sequencing (ChIP-seq), DNase I hypersensitive sites sequencing (DNase-seq), global run-on sequencing (GRO-seq), cap analysis gene expression (CAGE), etc. (11-16). In ChIP-seq experiments, genomic regions enriched with H3K4me1 and H3K27ac modifications are widely considered as active enhancers, and those with H3K4me1 and H3K27me3 modifications are taken as repressed enhancers (13). In DNase-seq, distal open chromatin regions are considered as potential enhancers for gene regulation studies (17-20). In GRO-seq and CAGE experiments, bidirectional transcripts are employed to identify active enhancers under specific experimental conditions (14,21,22). Correspondingly, computational methods based on NGS data are developed to predict enhancers on the genome-wide scale (13,23-25). These methods range from the early ones that are based solely on H3K4me3 and H3K4me1 ChIP-seq experiments to the later ones that are based on various types of epigenomic and genomic signals and have been applied to predict enhancers in different cell lines.

A large number of enhancers have been discovered so far. For instance, about 2,900 enhancers from comparative genomics were tested with mouse transgenic reporter assay and stored in the VISTA database (26). The Functional Annotation of the Mouse/Mammalian Genome (FANTOM) project (http://FANTOM.gsc.riken.jp/5/datafiles/latest/extra/Enhancers/) identified 32,693 enhancers from balanced bidirectional capped transcripts(14). This set of enhancers is arguably the largest set of mammalian enhancers with supporting experimental evidence (27). There are also hundreds of thousand computationally predicted human enhancers, such as those predicted by ChromHMM and Segway (23,24). This set of enhancers represents the most comprehensive set of computationally predicted human enhancers currently available although they are much less reliable.

Despite this relatively effortless discovery of enhancers, the identification of IEPs is still nontrivial. Early experimental procedures to identify IEPs are expensive and time-consuming (28,29). Recent Hi-C experiments hold a great promise to identify IEPs on the genome-scale, while are still are not cost effective in order to generate high-resolution Hi-C interactions (30-32). To date, these experiments have only been carried out on a few cell lines or cell types. Although computational methods, from the early ones defining the closest genes as target genes, to the later ones considering the correlation of epigenomic signals in enhancers and those in promoters, to the current ones based on more sophisticated approaches (14,18,33-39), have shown some success in predicting enhancer target genes, they either do not consider or have low–performance on cell-specific IEP prediction (35, 48).

All existing computational methods almost always consider one enhancer-promoter pair at a time to determine whether they interact. We hypothesized that when two enhancers interact with a common target gene, these two enhancers may be spatially close to each other and may thus interact with all target genes of both enhancers. In other words, if two enhancers share a target gene, they may as well share all of their target genes. If this hypothesis is true, we should consider the interactions of multiple enhancers and multiple target genes simultaneously to predict IEPs, which may improve the accuracy of the computational prediction of the IEPs, especially that of cell-specific IEPs.

To test this hypothesis, we collected IEPs determined in four previous studies (4,30-32) and investigated how different enhancers may share target genes under specific experimental conditions (Material and Methods). We observed that two enhancers are likely to either share almost all of their target genes or interact with two completely disjoint sets of target genes, under a given experimental condition. This observation implies an interesting characteristic of IEPs, which has not been considered by existing studies to predict IEPs. Our study may also shed new light on the underlying principles of chromatin interactions and facilitate the more accurate identification of IEPs.

## RESULTS

### Two enhancers are likely to interact with either exactly the same set or two completely different sets of genes

In order to study IEPs, we calculated the bipartite clustering coefficient (BCC) of enhancers in each cell line or cell type, with two sets of enhancers and four sets of IEPs (Material and Methods, Figure 1A). BCC is commonly used in a bipartite graph to measure how nodes share their neighboring nodes. For a given set of IEPs, its enhancer set and its gene set correspond to the two disjoint sets of nodes, and their interactions correspond to edges in a bipartite graph (Figure 1B). The neighboring nodes of an enhancer are the target genes of this enhancer. We thus can use the BCC to measure the percentage of target genes a pair of enhancers may share in this set of IEPs (Figure 1B). We observed that the BCC of enhancers was usually larger than 0.90. This indicates that when two enhancers interact with a common target gene, both enhancers are likely to interact with all target genes of these two enhancers.

**Figure 1:**
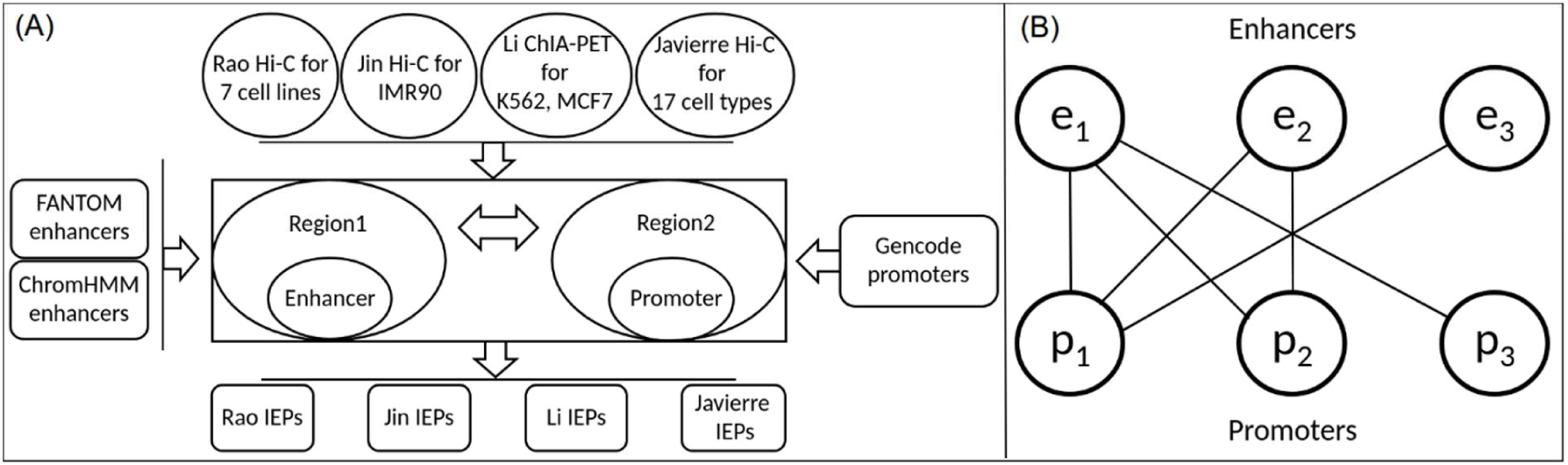
**(A)** The process of generating IEPs using the chromatin interaction data of Rao et al., Jin et al., Li et al., and Javierre et al., enhancer regions from FANTOM and ChromHMM and promoters defined around the GENCODE annotated gene TSSs. **(B)** A toy interaction network between three enhancers (*e*_1_, *e*_2_ and *e*_3_) and three promoters (*p*_1_, *p*_2_ and *p*_3_). Here, 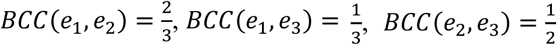 and 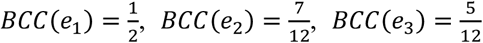. The average BCC of the enhancers in this example is 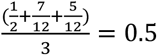.

First, we studied the IEPs based on the looplists from Rao et al. (31), with the annotated FANTOM enhancers (14) and the GENCODE promoters (41) (Figure 1A). We noticed that the BCC of enhancers was no smaller than 0.96 in all cell lines with enough IEPs (Table 1 and Supplementary Table S1). We further calculated the average BCC of the enhancers interacting with more than one gene. We found that their average BCC was no smaller than 0.94 in all cell lines, suggesting that two enhancers are likely to interact with either the same set or two disjoint sets of target genes.

**Table 1:**
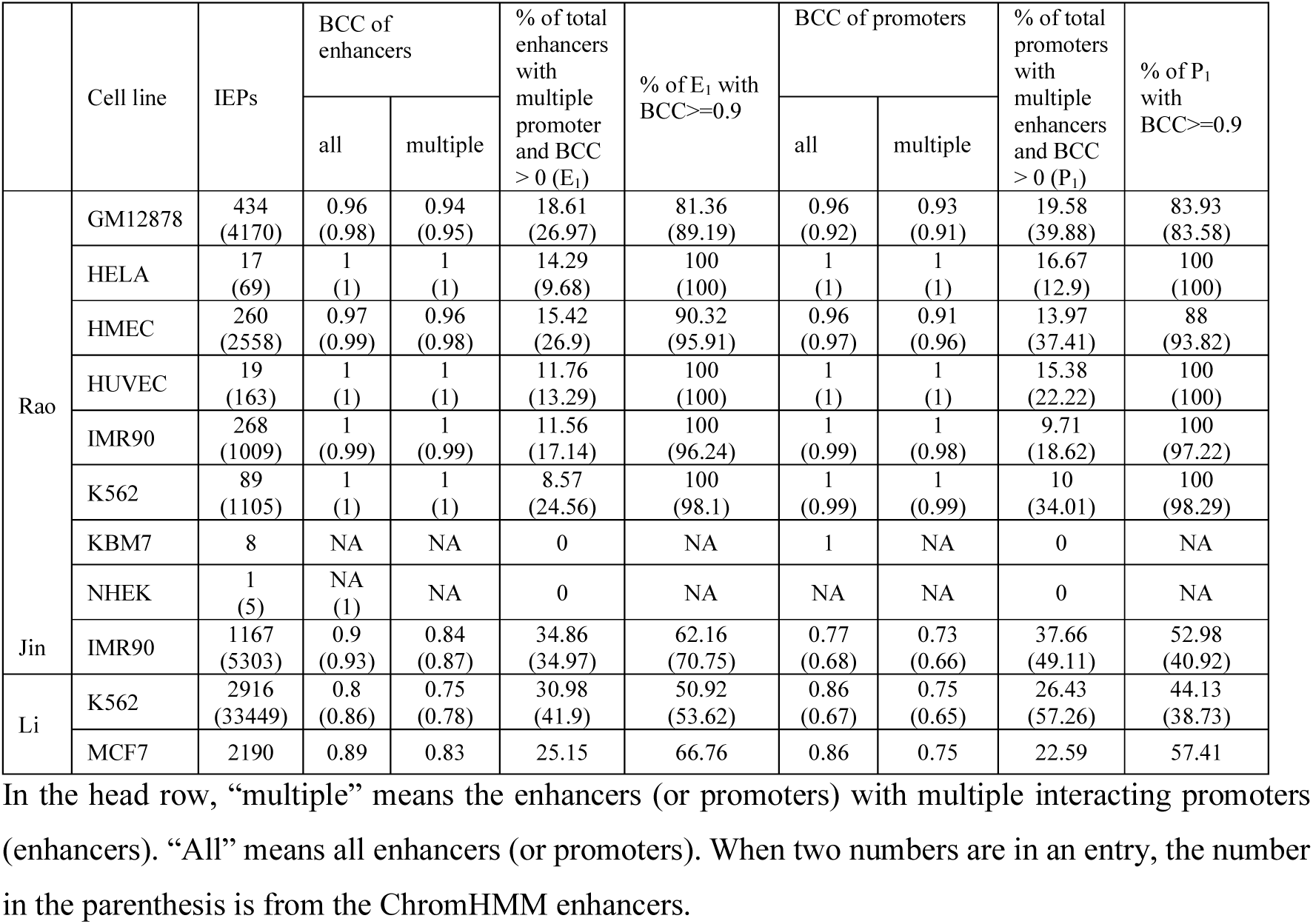
The BCC of enhancers and that of promoters are likely to be 1 under different experimental conditions.

| | Cell line | IEPs | BCC of enhancers | | % of total enhancers with multiple promoter and BCC > 0 ($E_1$) | % of $E_1$ with BCC >= 0.9 | BCC of promoters | | % of total promoters with multiple enhancers and BCC > 0 ($P_1$) | % of $P_1$ with BCC >= 0.9 |
| --- | --- | --- | --- | --- | --- | --- | --- | --- | --- | --- |
|  |  |  | all | multiple |  |  | all | multiple |  |  |
| Rao | GM12878 | 434<br>(4170) | 0.96<br>(0.98) | 0.94<br>(0.95) | 18.61<br>(26.97) | 81.36<br>(89.19) | 0.96<br>(0.92) | 0.93<br>(0.91) | 19.58<br>(39.88) | 83.93<br>(83.58) |
|  | HELA | 17<br>(69) | 1<br>(1) | 1<br>(1) | 14.29<br>(9.68) | 100<br>(100) | 1<br>(1) | 1<br>(1) | 16.67<br>(12.9) | 100<br>(100) |
|  | HMEC | 260<br>(2558) | 0.97<br>(0.99) | 0.96<br>(0.98) | 15.42<br>(26.9) | 90.32<br>(95.91) | 0.96<br>(0.97) | 0.91<br>(0.96) | 13.97<br>(37.41) | 88<br>(93.82) |
|  | HUVEC | 19<br>(163) | 1<br>(1) | 1<br>(1) | 11.76<br>(13.29) | 100<br>(100) | 1<br>(1) | 1<br>(1) | 15.38<br>(22.22) | 100<br>(100) |
|  | IMR90 | 268<br>(1009) | 1<br>(0.99) | 1<br>(0.99) | 11.56<br>(17.14) | 100<br>(96.24) | 1<br>(0.99) | 1<br>(0.98) | 9.71<br>(18.62) | 100<br>(97.22) |
|  | K562 | 89<br>(1105) | 1<br>(1) | 1<br>(1) | 8.57<br>(24.56) | 100<br>(98.1) | 1<br>(0.99) | 1<br>(0.99) | 10<br>(34.01) | 100<br>(98.29) |
|  | KBM7 | 8 | NA | NA | 0 | NA | 1 | NA | 0 | NA |
|  | NHEK | 1<br>(5) | NA<br>(1) | NA | 0 | NA | NA | NA | 0 | NA |
| Jin | IMR90 | 1167<br>(5303) | 0.9<br>(0.93) | 0.84<br>(0.87) | 34.86<br>(34.97) | 62.16<br>(70.75) | 0.77<br>(0.68) | 0.73<br>(0.66) | 37.66<br>(49.11) | 52.98<br>(40.92) |
| Li | K562 | 2916<br>(33449) | 0.8<br>(0.86) | 0.75<br>(0.78) | 30.98<br>(41.9) | 50.92<br>(53.62) | 0.86<br>(0.67) | 0.75<br>(0.65) | 26.43<br>(57.26) | 44.13<br>(38.73) |
|  | MCF7 | 2190 | 0.89 | 0.83 | 25.15 | 66.76 | 0.86 | 0.75 | 22.59 | 57.41 |

In the head row, “multiple” means the enhancers (or promoters) with multiple interacting promoters (enhancers). “All” means all enhancers (or promoters). When two numbers are in an entry, the number in the parenthesis is from the ChromHMM enhancers.

To assess the statistical significance of the above observation, we studied the BCC of enhancers in randomly generated IEPs (Supplementary Table S2). These random lEPs were constructed using the same set of enhancers and promoters but randomized interactions, where we randomly chose promoters to interact with an enhancer so that the same enhancer had the same number of interactions as it had originally. We also made sure that the random IEPs have a minimum distance of 2.5 kilobase pairs (kbps) as we did for the original IEPs. We generated five different sets of random IEPs in this way with five different random seeds. With these random IEPs in the eight cell lines, we barely had a handful of enhancers sharing promoters with other enhancers in any cell line, suggesting that it is not by chance that multiple enhancers interact with a common set of target genes in the Rao et al.’s looplists (Supplementary Tables S1 and S2). For all three cell lines we could calculate the BCC, the BCC of enhancers was 0.20, 0.33, and 0.62, respectively, which was much smaller than the BCC of enhancers in the above sets of “real” IEPs. When we considered the BCC of enhancers interacting with multiple genes, it was no larger than 0.17 for random IEPs, while it was no smaller than 0.94 for the “real” IEPs, also suggesting that the observation that the BCC of enhancers being close to 1 was not by chance.

Second, we studied IEPs defined by different cutoffs in seven cell lines (Material and Methods). Compared with the IEPs from the above Rao et al.’s looplists, these IEPs defined by cutoffs were likely to include many more bona fide IEPs and more false positives as well. Under the cutoffs 30, 50 and 100, the BCC of enhancers in all seven cell lines except GM12878 was no smaller than 0.73, 0.84 and 0.91, respectively (Supplementary Table S1). Since GM12878 had a much higher sequencing depth than other cell lines, it was understandable that a stringent cutoff for other cell lines was still loose for GM12878. We thus tried the cutoffs 150, 200, 300, and 400 for GM12878. We noticed that the BCC of enhancers was 0.93 in GM12878 with the cutoff 400. Coincidently, the number of IEPs in GM12878 defined at the cutoff 400 was similar to that in other cell lines defined at the cutoff 100 (Supplementary Table S1). Note that in HMEC, HUVEC, KBM7 and NHEK, the BCC of enhancers was no smaller than 0.90 even under the cutoff 100. Moreover, the BCC of enhancers was increasing with more stringently defined IEPs, suggesting that the BCC of enhancers is close to 1 if it is not 1 (Supplementary Table S1).

In order to assess the statistical significance of the observed BCC of enhancers in IEPs from different cutoffs, we similarly generated random IEPs (Supplementary Table S2). Again, for every cutoff in every cell line, the BCC of enhancers for random IEPs was much smaller than the BCC of enhancers for “real” IEPs. For instance, under the cutoff 30, the BCC of enhancers was 0.33 and 0.34 for random IEPs in IMR90 and K562, respectively, while the corresponding number was 0.74 and 0.73 for “real” IEPs. If we considered only enhancers interacting with multiple target genes, the BCC of enhancers for random IEPs was about two times smaller than that for “real” IEPs. For instance, under the cutoff 30, such a BCC was 0.25 and 0.25 for random IEPs in IMR90 and K562, respectively, while the corresponding number was 0.66 and 0.64 for “real” IEPs, respectively.

Third, to see how this observation might change if we used the data from other labs or other experimental protocols, we studied the IEPs from three additional studies (Figure 1A, Material and Methods) (4,30,32). When we calculated the BCC of enhancers using the IEPs defined by Jin et al. themselves (30), it was 0.94. When considering the IEPs defined by Jin et al. based on the annotated enhancers by FANTOM and the annotated promoters by GENCODE, it was 0.90. In terms of the ChIA-PET datasets (4), it was 0.80 in K562 and 0.89 in MCF7. For the nine cell types from Javierre et al. (32), it was no smaller than 0.96 in all cell types (Supplementary Table S1). Although the IEPs were from different labs and from different experimental procedures, in all cases, the BCC of enhancers was larger than 0.80 and the majority of enhancers interacting with multiple promoters had their individual BCCs larger than 0.90, suggesting that the BCC of enhancers is likely to be 1 under an experimental condition. Again, for the corresponding randomly generated IEPs for these datasets, it was 0.30 on average in all cases, much smaller than the corresponding ones from original IEPs, which was 0.96 on average (Supplementary Table S2).

Finally, we repeated the above analyses with the FANTOM enhancers replaced by the ChromHMM enhancers (23). We had a similar observation in all cases (Supplementary Table S1). That is, the BCC of enhancers for IEPs in a cell line was close to 1. For instance, for IEPs based on the looplists, it was 1 or no smaller than 0.98 in all cell lines. For the Hi-C data from Rao et al. under the cutoff 100, it was no smaller than 0.90 in all cell lines. For the Hi-C data from Jin et al. (6), it was 0.93. For the ChIA-PET data from Li et al. (30), it was 0.86. For the nine cell types from Javierre et al. (32), it was no smaller than 0.97. In almost all cases, the majority of enhancers with multiple promoters had their individual BCCs larger than 0.90.

In summary, the BCC of enhancers was likely to be close to 1 for the different set of IEPs generated with the data from different labs, different experiments, different cell lines and cell types, and different enhancer sets. The analyses based on IEPs from different cutoffs suggest that the BCC of enhancers is quite robust, although it is smaller when more loosely defined IEPs are used. It is close to 1 or becomes 1 when the IEPs are defined more and more stringently (with fewer false positive IEPs). These analyses suggest that what we observed may be an intrinsic property of enhancers. That is, if two enhancers interact with one common gene, they are likely to interact with each of their individual target genes.

### Two target genes tend to interact with exactly the same set or two completely different sets of enhancers

We studied the BCC of promoters in each set of the aforementioned IEPs to see whether the similar hypothesis was true for the BCC of promoters. Our data showed that the BCC of promoters was likely to be 1 as well, although this was not so evident as the BCC of enhancers in certain cases.

First, we studied the BCC of promoters with IEPs based on the looplists (31). It was close to 1 no matter whether we used the FANTOM enhancers or the ChromHMM enhancers (Table 1). We also calculated the BCC of promoters in randomly simulated IEP datasets, where we kept the same sets of enhancers and promoters but randomly selected enhancers to interact with a promoter so that every promoter had the same number of interacting enhancers as it had in the original set of IEPs. The BCC of promoters was not larger than 0.53 in any cell line in these random datasets, suggesting that it was not by chance that the BCC of promoters was close to 1 in all cell lines (Supplementary Tables S1 and S2).

Second, we studied the BCC of promoters based on lEPs defined with different cutoffs (31) (Supplementary Table S1). When we used the FANTOM enhancers, the BCC of promoters was often close to 1. For instance, with the cutoff 400 for GM12878 and the cutoff 100 for other cell lines, the BCC of promoters was no smaller than 0.90 in all cell lines. For different cutoffs, it was usually no smaller than the BCC of enhancers, which was close to 1 in most cases. When we used the ChromHMM enhancers, however, it was not as large as those from the FANTOM enhancers. For instance, with the cutoff 400 for GM12878 and the cutoff 100 for other cell lines, the BCC of promoters varied from 0.66 to 0.87 in different cell lines. It was smaller when the cutoff was smaller. This might be due to the much lower quality of the ChromHMM enhancers compared with the experimentally defined FANTOM ones.

Although the BCC of promoters was not as large as the BCC of enhancers when the ChromHMM enhancers were used, the actual BCC of promoters could also be close to 1. This was because the computationally predicted ChromHMM enhancers might result in predicting false interactions and thus a low BCC of promoters. Moreover, the BCC of promoters was always increasing with more and more stringently defined IEPs. Although we did not observe that the BCC of promoters was close to 1 at the cutoff 100 we tried, it was indeed close to 1 when the looplists defined by Rao et al. were considered. In addition, the BCC of promoters for random IEPs in every cell line and under every cutoff was much smaller than that for the “real” IEPs, indicating that the observed much larger BCC of promoters was not by chance (Supplementary Table S2).

Third, we studied the BCC of promoters based on lEPs from other studies (Figure 1B, Table 1 and Supplementary Table S1) (4,30,32). For the original IEPs from Jin et al., it was 0.11. However, when the IEPs were defined from the overlap of these original IEPs with the GENCODE promoters and the two types of enhancers, it was 0.77 and 0.68, respectively (Table 1). The low BCC of promoters for the original IEPs may be partially due to the promoters Jin et al. used, which had 11,313 promoters inferred by Jin et al., compared to the 57,820 promoters annotated by GENCODE. In terms of the ChIA-PET data, when we used the FANTOM enhancers, the BCC of promoters was 0.86 in K562 and 0.86 in MCF7; when we used the ChromHMM enhancers, it was 0.67 in K562. ChromHMM did not have annotated enhancers in MCF7. For the nine cell types from Javierre et al., it was no smaller than 0.98 and 0.91 when the FANTOM enhancers and the ChromHMM enhancers were used, respectively. Overall, although it was not as large as the BCC of enhancers, because of the imperfectness of all these collected IEPs, and the fact that the majority of promoters interacting with multiple enhancers had their individual BCC larger than 0.90, and they were much larger than the corresponding BCC of promoters for random IEPs (0.30), the BCC of promoters was likely to be close to 1 as well. In other words, two target genes likely interact with the same set of enhancers or two disjoint sets of enhancers.

### Enhancers form clusters that have special characteristics

Since the BCC of enhancers is close to 1, we can organize enhancers into clusters, where every enhancer in the same cluster is likely to interact wtih the same set of target genes. We thus built an enhancer graph by connecting enhancers that share at least one common target. We then grouped enhancers into clusters based on such a graph in each cell line (Material and Methods, Figure 2). Here we only considered the looplists and the IEPs obtained from the most stringent cutoff (400 in GM12878 and 100 in other cell lines) to obtain enhancer clusters, as they were more reliable than other sets of IEPs.

**Figure 2:**
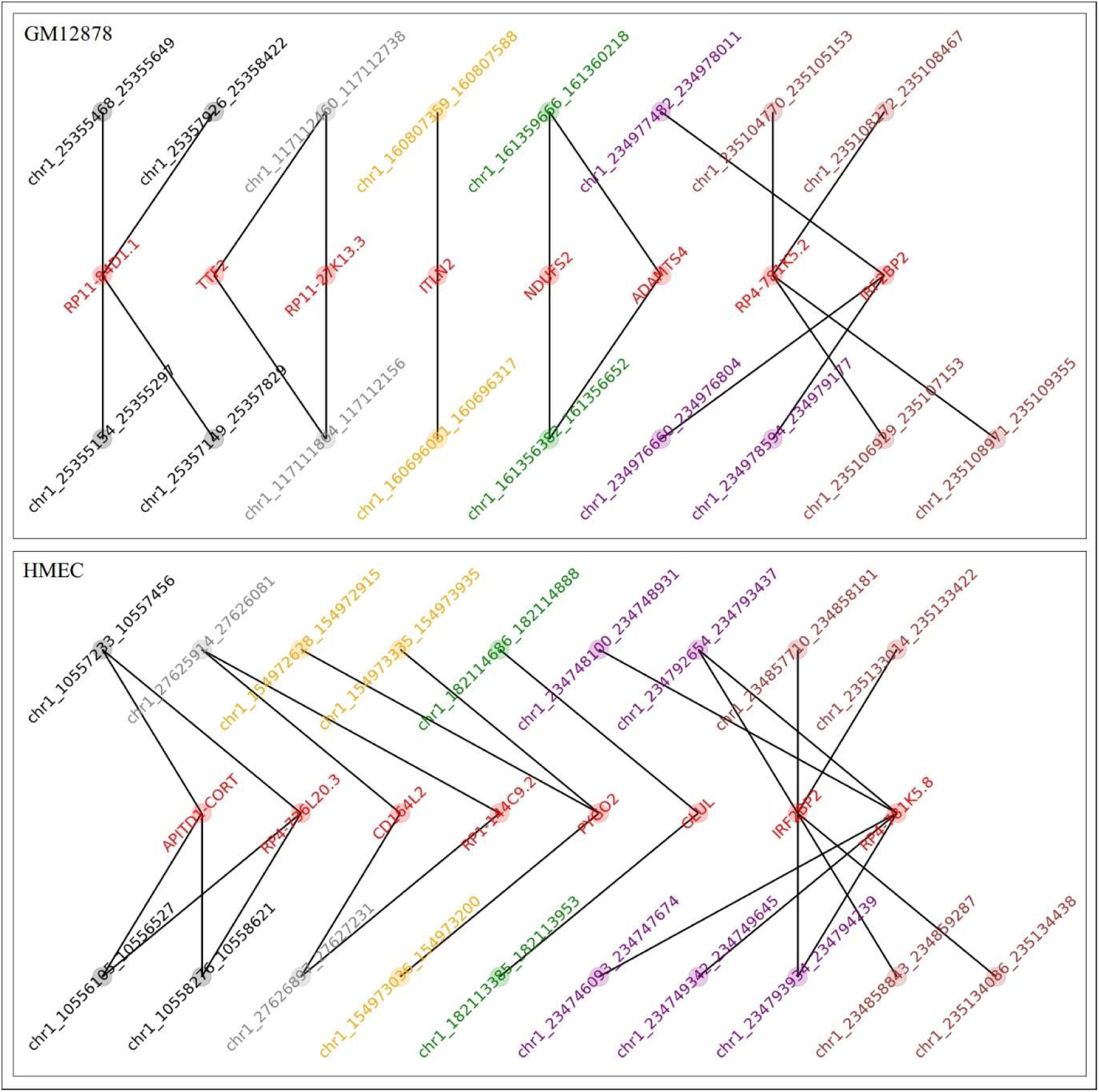
Cluster of enhancers (nodes in the top and bottom rows in each connecting subgraph) interacting with the same genes (nodes in the middle row) in chromosome 1 in GM12878 and HMEC. The clusters were defined from the IEPs found using Rao et al. Hi-C looplist data. The enhancers and the gene nodes are arranged from left to right according to their relative genomic locations.

We obtained 1 to 2,755 clusters in different cell lines. The number of clusters in a cell line and across different cell lines varied dramatically, depending on the IEPs and the enhancers used (Supplementary Table S3). When the ChromHMM enhancers were used, there were many more clusters and 60% to 96% of all enhancers were included in clusters. When the FANTOM enhancers were used, fewer clusters were identified and about 14% to 59% of the total enhancers were in clusters. The average number of enhancers in a cluster varied from 2 to 5 in different cell lines. Enhancers in the majority of clusters interacted with only one gene, while on average, enhancers in 22.91% clusters interacted with at least two different genes.

We studied the distance between the enhancers in a cluster and the distance between enhancers and their target genes (Figure 3 and Supplementary Table S4). On average, about 96% of the enhancers in a cluster were within 10 kbps. However, there was a small fraction of enhancers in a cluster that were more than 50 kbps away from each other. For instance, when the looplists and the FANTOM enhancers were considered, there were more than 8% enhancers in a cluster that were more than 50 kbps away from each other in GM12878, HMEC and IMR90. In terms of the target genes, the majority of them were within 10 kbps, with a small fraction far from each other. For instance, when the looplists and the FANTOM enhancers were considered, we found 26.92%, 21.43% and 7.41% of the target genes of an enhancer cluster that were more than 50 kbps away from each other in GM12878, HMEC and IMR90, respectively. It was also worth pointing out that the enhancers in a cluster were normally consecutive active enhancers while their target genes were normally not consecutive. In all cell lines, on average, more than 93% of the enhancers in a cluster were consecutive active enhancers while fewer than 22% of the target genes of an enhancer cluster were consecutive.

**Figure 3:**
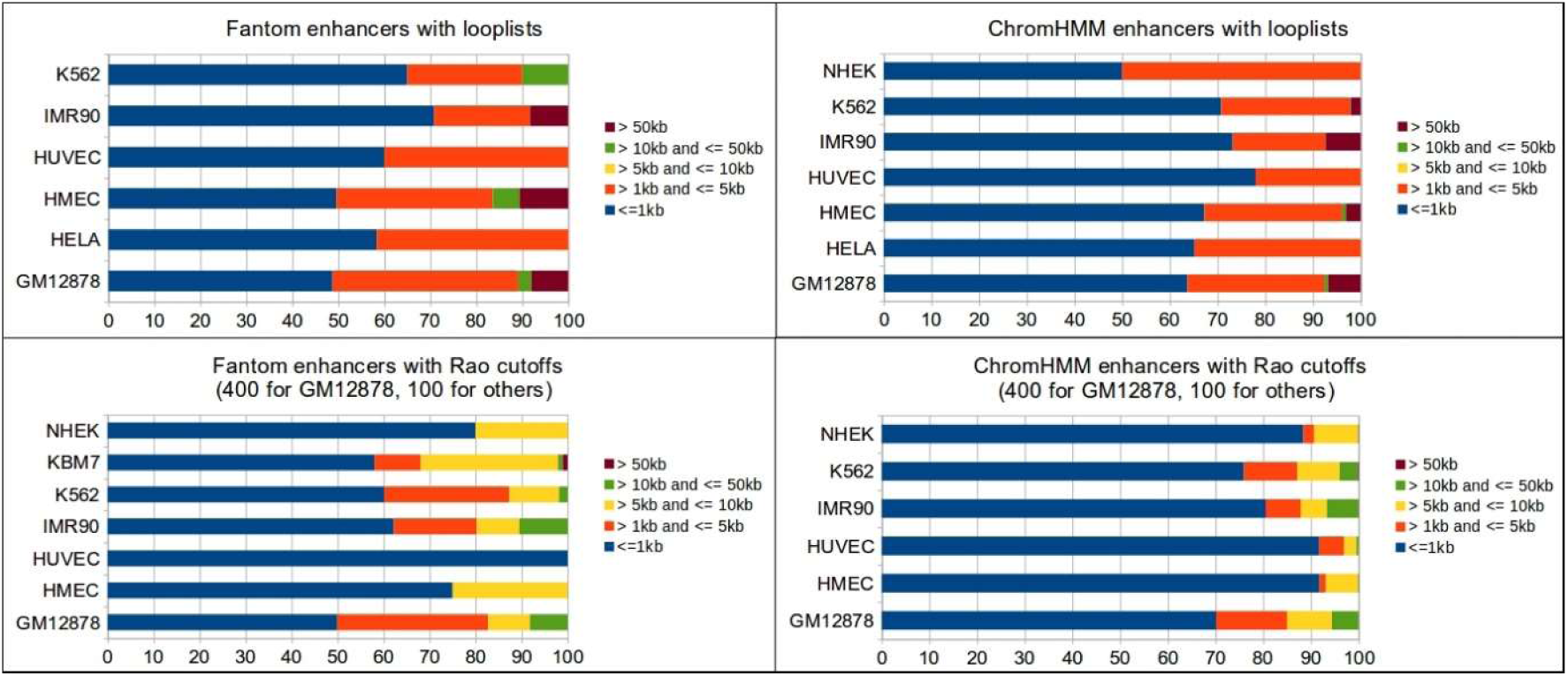
Average distance distribution between enhancers in the same cluster for each cell line. The X-axis represents the average percentage of enhancers in each enhancer cluster maintaining a given distance range from their nearest enhancers in that cluster.

Since enhancers in a cluster were consecutive in the genome and the majority of enhancers in a cluster were close to each other, they seemed like the super-enhancers. We thus compared the enhancer clusters with known super-enhancers (Supplementary Table S5). On average, 26.70% of enhancer clusters overlapped with the corresponding super-enhancers in a cell line while the majority of enhancer clusters did not overlap with the known super-enhancers (Figure 4A), which may represent new super-enhancers. On the other hand, a large proportion of known super-enhancers did not overlap with the enhancer clusters in the corresponding cell lines (Figure 4B). Interestingly, when a super-enhancer overlapped an enhancer cluster, more than 75% of the genomic regions that contain all enhancers in this enhancer cluster were within this super-enhancer.

**Figure 4:**
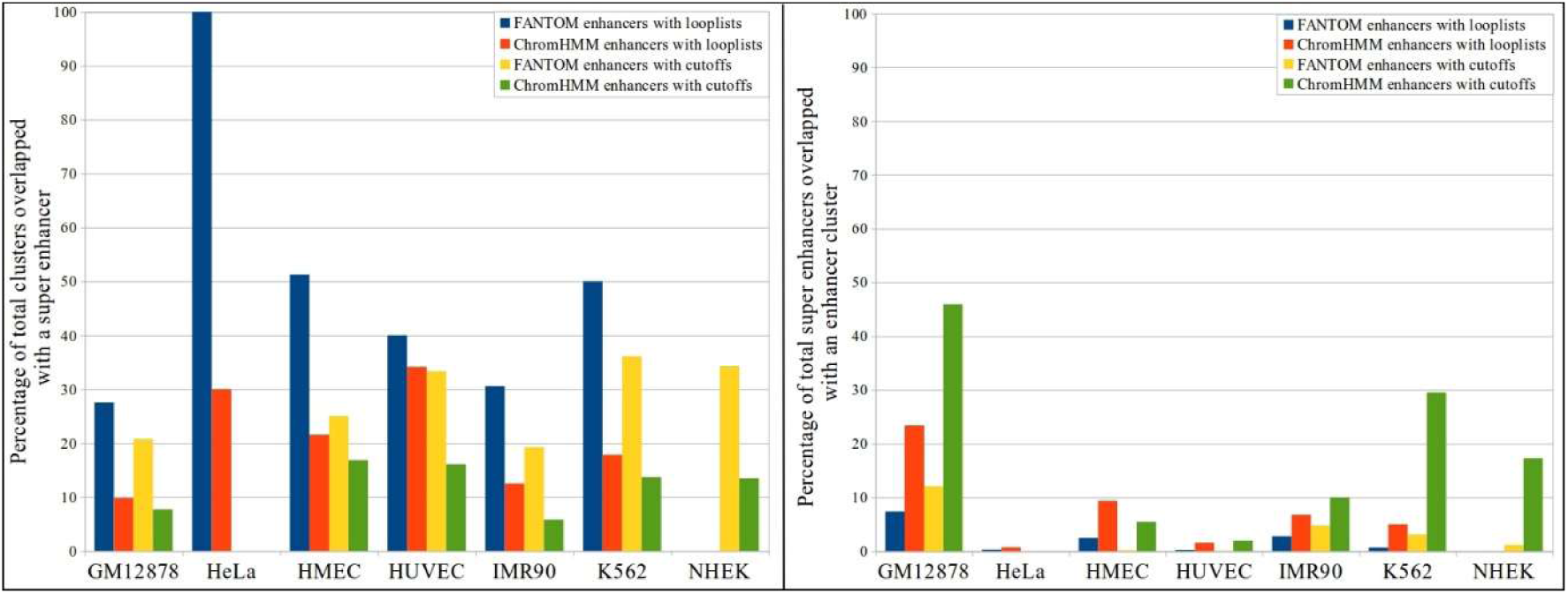
The overlap of the enhancer clusters with the super-enhancers. **(A)** The percentage of the enhancer clusters overlapping with the super-enhancers. **(B)** The percentage of the super-enhancers overlapping with the enhancer clusters.

We also studied how the enhancers in a cluster located relative to a topologically associating domain (TAD) (Supplementary Table S6). The enhancers in a cluster were usually within the same TAD, with no smaller than 98.38% of enhancers in a cluster within a TAD in every cell line, independent of IEPs and enhancers used. In most cell lines, for all clusters, all enhancers in a cluster were within a TAD. The slight deviation from the 100% in certain cases may be due to the imperfectness of the IEPs, enhancers, and TADs, mostly due to the computationally predicted enhancers, as the percentage was always 100% in all cell lines except GM12878 when the FANTOM enhancers were used.

We studied how the enhancer clusters were shared by different cell lines as well (Supplementary Table S7). That is, for an enhancer cluster in a cell line, how likely was the same cluster identified in another cell line. We found that on average no more than 14% enhancer clusters were identified in two cell lines. Moreover, the percentage was smaller when the looplists were used than when the stringent cutoffs were used to define IEPs, implying that the looplists were too strict to include many bona fide IEPs. The small percentage of shared enhancer clusters suggested that most enhancer clusters were cell-specific.

## DISCUSSION

We observed that two enhancers either do not share any target gene or share almost all of their target genes. This observation was true when different sets of IEPs, two sets of enhancers, and a variety of cell lines and cell types were considered. Moreover, the BCC of enhancers became closer and closer to 1 when the criteria to define IEPs became more and more stringent. In addition, the same observation did not hold to be true for randomly generated IEPs. These analyses suggested that the BCC of enhancers under a particular experimental condition was likely to be close to 1 if it is not 1.

Similarly, we observed that two promoters were likely to interact with either the same set of enhancers or two disjoint sets of enhancers. This observation about promoters was not as evident as that about enhancers. However, it was pervasive in all cases when the FANTOM enhancers were used. It was also evident when the looplists and the IEPs defined by the most stringent cutoffs were used. Although it seemed not compelling when the ChromHMM enhancers and the sets of IEPs that were defined with loose criteria were used, this might be due to the imperfectness of enhancers and IEPs we had. More importantly, the fact that the BCC of enhancers was close to 1 implied that the BCC of the promoters should be close to 1 as well based on the definition of the BCC.

The BCC of enhancers being close to 1 suggested that enhancers form clusters to interact with the target genes. As shown above, these clusters are different from the known enhancer clusters such as super-enhancers, although they do overlap in certain regions. Enhancers in the clusters here were likely to interact with the same set of genes, while enhancers in a super-enhancer do not necessarily interact with a common target gene. Moreover, the enhancers in a cluster here could be far from each other while the enhancers in a super-enhancer are quite close to each other.

The BCC of enhancers was not 1 sometimes, which implied that when a group of enhancers interacts with a set of target genes, the majority of target genes interact with each enhancer in this group while the rest interact with only a subset of enhancers in this group. We called the former the fully shared target genes and the latter the partially shared target genes. The percentage of the partially shared target genes by a group of enhancers varied from 0% to 9.48%. We compared these two types of target genes in terms of TAD, tissue specificity, and correlations with the enhancers, with the IEPs from the looplists and the IEPs from the most stringent cutoff (400 in GM12878 and 100 in other cell lines) (Material and Methods). We did not observe any difference between the two types of target genes.

In practice, several aspects may prevent the BCC of enhancers and the BCC of promoters from being 1. First, the resolution of the interaction data prevents from obtaining accurate IEPs. The two interacting regions in the interaction data are often long, which is around 5 kbps in most of the cases we studied. We defined IEPs by overlapping enhancers and promoters with pairs of interacting regions, which might be prone to errors, given the fact that many known enhancers were much shorter (46, 47). Second, the IEPs defined imperfectly might have produced “false” interactions and thus decreased the BCCs. Third, the enhancers were not perfectly defined either. The FANTOM enhancers are still far from complete while the computationally predicted ChromHMM enhancers may contain many “false” enhancers.

Despite the limitations, it was evident that under a particular experimental condition, the BCC of enhancers and the BCC of promoters are likely to be close to 1. This suggests that active enhancers form clusters to interact with a group of genes. Similarly, active promoters form clusters that interact with the same group of enhancers. It is thus important to consider the relationship among enhancers and among promoters when studying their interactions, which may help improve our understanding of the distal gene regulation and the chromatin structures.

## MATERIAL AND METHODS

### Enhancers and promoters

To study IEPs, we considered two sets of enhancers (Figure 1B). The first set contained the 32,693 enhancers annotated by FANTOM, which had been obtained from the balanced bidirectional capped transcripts in CAGE experiments (14). The second set was the computationally predicted enhancers by ChromHMM (23) in the following seven cell lines: GM12878, HMEC, HUVEC, K562, NHEK, IMR90 and HeLa. ChromHMM is a widely used tool to partition genomes into different functional units including enhancers.

The FANTOM enhancers are not cell-specific, while the ChromHMM predicted enhancers are specific for the seven different cell lines mentioned. We thus defined cell-specific “active” FANTOM enhancers, by overlapping the enhancers with the H3K27ac ChIP-seq peaks in the corresponding cell lines obtained from the Encyclopedia of DNA Elements (ENCODE) project (40). When there was no H3K27ac ChIP-seq data available for a cell line such as KBM7, we considered the enhancers that overlapped with the chromatin interacting anchors in this cell line as “active” enhancers (31).

We used the gene transcriptional start sites annotated in the GENCODE V19 (41) to define promoters. The upstream region of 1 kbps to the downstream region of 100 bps of each transcriptional start site was considered as a promoter. In total, we obtained 57,820 promoters in the human genome.

### IEPs from four previous studies

To learn the new characteristics of IEPs, we collected IEPs from four previous studies (4,30-32) (Figure 1B). These IEPs arguably represent the chromatin interactions defined with the highest resolutions by the corresponding techniques (4,30-32). The first set of IEPs was from the Hi-C dataset GSE63525 in the Gene Expression Omnibus (GEO) database, where Rao et al. extracted significant intra-chromosomal chromatin interactions called “looplists” in the following eight cell lines: GM12878, HeLa, HMEC, HUVEC, IMR90, K562, KBM7 and NHEK (31). These looplists were defined with stringent criteria and were most likely to be true pairs of interacting genomic regions, each of which was about 5 kbps long (Supplementary Table S1). In every cell line, we overlapped each looplist with the aforementioned two sets of enhancers and with the annotated promoters to obtain IEPs. In other words, an obtained IEP consisted of an enhancer and a promoter, where the enhancer overlapped with one region specified in a looplist and the promoter overlapped with the other region specified in the same looplist. Since we had two sets of enhancers, we obtained two sets of IEPs for each of the eight cell lines (Figure 1A).

Note that the number of IEPs obtained from the above looplists was small, especially when we considered the FANTOM enhancers (Supplementary Table S1). The reason might be, the criteria Rao et al. used to define looplists was quite stringent and many true interacting genomic regions might therefore be missed (48). To capture more IEPs in these cell lines, we also defined alternative sets of IEPs with three cutoffs: 30, 50, and 100, from the normalized Hi-C contact matrices defined by Rao et al. (31). Given a normalized read cutoff, say x, if an enhancer-promoter pair overlapped with a pair of interacting genomic regions that were supported by at least x normalized Hi-C reads, we considered this EP-pair as an IEP. The cutoff 30 was used since this cutoff was likely to include of almost all known IEPs in K562 and IMR90 from other studies (6, 30) without allowing too many false positives (48). The two other cutoffs were used to see how the observed enhancer characteristics may change with more stringent cutoffs. Intuitively, the larger the cutoff was, the more likely that the two genomic regions interacted. Based on our previous studies (37,42), we believed that the majority of the IEPs defined by these cutoffs in the eight cell lines except HeLa and GM12878 were likely to be bona fide IEPs and considered the IEPs defined by the cutoff 100 as highly reliable. We could not define IEPs in HeLa by cutoffs because Rao et al. did not provide a Hi-C contact matrix in HeLa. Since the sequencing depth was much higher in case of GM12878 than that in other seven cell lines, we considered the IEPs defined by the cutoff 400 in GM12878 as highly reliable after testing different cutoffs.

We also obtained 57,578 IEPs in IMR90 from another Hi-C study (30). To our knowledge, this was the only Hi-C dataset for human samples with a comparable sequencing depth as that in GSE63525. In this study, Jin et al. defined active enhancers with H3K4me1 and H3K27ac ChIP-seq peaks and active promoters with H3K4me3 ChIP-seq peaks together with the known genes from the University of California, Santa Cruz genome browser. In addition to using the original IEP dataset which was provided in the hg18 version (30), we also converted it into the hg19 version and overlapped with the aforementioned enhancers and promoters used in this study to define IEPs.

We used the IEPs defined by the ChIA-PET experiments in K562 and MCF7 as well for this study (4). ChIA-PET datasets for other cell lines have much lower sequencing depth than the two cell lines used here. Using the interacting regions in these datasets we found 2,923 and 2,190 IEPs with the FANTOM enhancers for K562 and MCF7, respectively. When we considered the ChromHMM enhancers, there were 33,598 IEPs for K562. There were no ChromHMM enhancers available in MCF7.

Finally, we obtained additional IEPs based on active enhancer and promoter links defined by Javierre et al. from promoter capture Hi-C experiments in nine cell types (Table S1 in (32)). Javierre et al. did the experiments on seventeen primary cell types while the active enhancer and promoter links were provided for nine cell types. Each link defined a pair of interacting regions, with the average length of 5,709 and 8,599 bps, respectively. Since Javierre et al. did not explicitly specify the enhancers and promoters, we overlapped these links with the two sets of enhancers and the GENCODE promoters to define two sets of IEPs. In total, we obtained 20,764 and 607,274 IEPs with FANTOM and ChromHMM enhancers, respectively.

We applied a distance filter on all the IEP sets found above. For every IEP, if the distance between the corresponding enhancer and promoter is less than 2.5kb, we filtered that IEP out from our analysis. The number of filtered IEPs for all the datasets are shown in Supplementary Table S1.

### Other data used

Rao et al. annotated chromatin contact domains in each of the eight cell lines (31). We downloaded these domains from GSE63525 and considered them as the topologically associating domains (TAD)s. We also downloaded the annotated TADs in IMR90 by Dixon et al., which were generated by the same lab that generated the Jin et al. data (43).

We downloaded the DNase I hypersensitive sites in the following thirteen cell lines: A549, CD14, CD20, CD20+RO01778, GM12878, H1, HeLa, HepG2, HUVEC, IMR90, K562, MCF7 and SKNSH from ENCODE (40). These are all ENCODE Tiers I and Tier II cell lines with the DNase-seq data were used in our previous study (37). For every IEP in each of the eight cell lines in GSE63525, we calculated the correlation of the DNase-seq signal in the enhancer region with the DNase-seq signal in the promoter region of the target gene across these thirteen cell lines. These correlations were used to compare different types of target genes based on the Mann-Whitney test.

We downloaded the super-enhancers in GM12878, HeLa, HMEC, HUVEC, K562 and NHEK from http://asntech.org/dbsuper/download.php. We could not find the known super-enhancers in KBM7. The super-enhancers in a cell line were compared with the clusters of enhancers that interact with the same set of target genes in the same cell line identified in this study.

We downloaded the CAGE peaks in 1,829 tissue samples from http://FANTOM.gsc.riken.jp/5/data/. By overlapping the GENCODE promoters with the CAGE peaks, we found the tissues in which the GENCODE genes are expressed. For each gene included in the defined IEPs, we recorded the list of the IDs of the tissues in which the gene is expressed. With this list of tissue IDs, we studied whether different types of target genes are expressed in the same tissues.

### BCC (Bipartite clustering coefficient)

All IEPs in a cell line form a bipartite graph, where the enhancers on one side connect with the target genes on the other side. We thus applied the BCC (44) to characterize how enhancers share their target genes and how genes share their enhancers (Figure 1A).

For a pair of enhancers (or a pair of genes), say *u* and *v*, their BCC is defined as 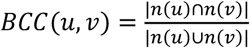, where *n(u)* and *n(v)* are the set of genes (or enhancers) interacting with *u* and *v*, respectively. Intuitively, if *u* and *v* are a pair of enhancers, *BCC(u,v)* measures the percentage of target genes both *u* and *v* interact with among all of their target genes. Similarly, if *u* and *v* are a pair of genes, *BCC(u,v)* measures the percentage of enhancers both *u* and *v* interact with among all enhancers they interact with. Correspondingly, the BCC of an individual enhancer (or gene), say *u*, is defined as 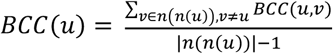, where *n(n(u))* is the set of enhancers (or genes) that share at least one target gene (or enhancer) with *u*. Under a given condition, for all enhancers (or target genes) sharing at least one target gene (or an enhancer) with other enhancers (or target genes), we averaged their individual BCCs to obtain the BCC of enhancers (or target genes) under this condition.

### Clusters of enhancers that interact with the common set of genes

We built an enhancer graph, with nodes representing enhancers and edges representing pairs of enhancers interacting with at least one common target gene. We applied the Bron-Kerbosch algorithm (45) to this graph to find all maximal cliques. Enhancers in a clique formed a cluster of enhancers that interact with the same set of genes. Different clusters may share enhancers.

## FUNDING

This work has been supported by the National Science Foundation [grants 1356524, 1661414 and 1149955] and the National Institute of Health [grant R15GM123407]. Funding for open access charge: The National Science Foundation grant 1149955.

## Notes

### Competing Interest Statement

The authors have declared no competing interest.

