## Supplementary Table S1-S7 for "An intriguing characteristic of enhancer-promoter interactions"

Table S1: BCC statistics for enhancers and promoters. The BCC of all enhancers (promoters) and the BCC of enhancers (promoters) interacting with multiple promoters (enhancers) are shown in the last four columns for different experimental conditions. ChromHMM Gencode Rao cutoff 30 could not be listed due to longer processing time.

| Experiments | Cell lines | IEPs | IEPs >= 2.5kb | Enhancers | Promoters | BCC of enhancers | | BCC of promoters | | Enhancers with BCC>0 | Enhancers interacting with multiple promoters and BCC>0 |
|---|---|---|---|---|---|---|---|---|---|---|---|
| | | | | | | all | with multiple targets | all | with multiple targets | | |
| FantomCage Gencode Rao looplist | GM12878 | 434 | 434 | 317 | 286 | 0.96 | 0.94 | 0.96 | 0.93 | 173 | 59 |
| | HELA | 17 | 17 | 14 | 12 | 1 | 1 | 1 | 1 | 7 | 3 |
| | HMEC | 260 | 260 | 201 | 179 | 0.97 | 0.96 | 0.96 | 0.91 | 102 | 31 |
| | HUVEC | 19 | 19 | 17 | 13 | 1 | 1 | 1 | 1 | 10 | 1 |
| | IMR90 | 268 | 268 | 199 | 206 | 1 | 1 | 1 | 1 | 83 | 24 |
| | K562 | 89 | 89 | 70 | 70 | 1 | 1 | 1 | 1 | 25 | 6 |
| | KBM7 | 8 | 8 | 5 | 8 | NA | NA | 1 | NA | 0 | 0 |
| | NHEK | 1 | 1 | 1 | 1 | NA | NA | NA | NA | 0 | 0 |
| FantomCage Gencode Rao cutoff 400 | GM12878 | 3822 | 1231 | 997 | 949 | 0.93 | 0.83 | 0.91 | 0.79 | 382 | 83 |
| FantomCage Gencode Rao cutoff 300 | GM12878 | 5527 | 2861 | 2138 | 2092 | 0.89 | 0.73 | 0.88 | 0.71 | 1012 | 234 |
| FantomCage Gencode Rao cutoff 200 | GM12878 | 8067 | 5240 | 3347 | 3328 | 0.81 | 0.69 | 0.82 | 0.68 | 2021 | 767 |
| FantomCage Gencode Rao cutoff 150 | GM12878 | 10072 | 6938 | 4179 | 4199 | 0.8 | 0.69 | 0.81 | 0.69 | 2614 | 1116 |
| FantomCage Gencode Rao cutoff 100 | GM12878 | 16391 | 13223 | 6080 | 6788 | 0.74 | 0.67 | 0.79 | 0.67 | 4424 | 2550 |
| | HMEC | 734 | 124 | 113 | 116 | 0.92 | 0.33 | 0.95 | 0.5 | 17 | 1 |
| | HUVEC | 179 | 31 | 30 | 28 | 1 | NA | 1 | NA | 6 | 0 |
| | IMR90 | 2870 | 1056 | 885 | 877 | 0.91 | 0.78 | 0.91 | 0.77 | 281 | 63 |
| | K562 | 2677 | 618 | 524 | 503 | 0.92 | 0.79 | 0.9 | 0.77 | 168 | 34 |
| | KBM7 | 7152 | 1204 | 1112 | 1093 | 0.95 | 0.69 | 0.94 | 0.71 | 204 | 17 |
| | NHEK | 2901 | 425 | 387 | 385 | 0.95 | 0.5 | 0.94 | 0.5 | 76 | 4 |
| FantomCage Gencode Rao cutoff 50 | GM12878 | 39558 | 36387 | 8906 | 13704 | 0.65 | 0.62 | 0.73 | 0.65 | 7552 | 6267 |
| | HMEC | 2896 | 511 | 453 | 456 | 0.95 | 0.8 | 0.95 | 0.83 | 92 | 15 |
| | HUVEC | 2422 | 454 | 401 | 392 | 0.92 | 0.72 | 0.9 | 0.69 | 103 | 15 |
| | IMR90 | 5589 | 3513 | 2455 | 2721 | 0.84 | 0.71 | 0.88 | 0.73 | 978 | 368 |
| | K562 | 5356 | 2820 | 2004 | 2101 | 0.85 | 0.72 | 0.87 | 0.71 | 878 | 281 |
| | KBM7 | 9191 | 2942 | 2489 | 2419 | 0.93 | 0.78 | 0.92 | 0.74 | 790 | 139 |
| | NHEK | 3348 | 780 | 672 | 659 | 0.92 | 0.74 | 0.91 | 0.72 | 183 | 33 |
| FantomCage Gencode Rao cutoff 30 | GM12878 | 70962 | 67791 | 9909 | 20636 | 0.6 | 0.59 | 0.68 | 0.62 | 9040 | 8422 |
| | HMEC | 4400 | 1718 | 1411 | 1380 | 0.93 | 0.8 | 0.92 | 0.76 | 485 | 98 |
| | HUVEC | 3910 | 1418 | 1148 | 1105 | 0.9 | 0.75 | 0.89 | 0.72 | 438 | 99 |
| | IMR90 | 10089 | 7951 | 4222 | 5348 | 0.74 | 0.66 | 0.84 | 0.69 | 2312 | 1309 |
| | K562 | 9138 | 6438 | 3458 | 4183 | 0.73 | 0.64 | 0.82 | 0.66 | 1985 | 1078 |
| | KBM7 | 13475 | 6608 | 5318 | 5043 | 0.9 | 0.76 | 0.88 | 0.71 | 2181 | 501 |
| | NHEK | 5304 | 2448 | 1974 | 1946 | 0.91 | 0.78 | 0.91 | 0.76 | 729 | 171 |
| ChromHMM Gencode Rao looplist | GM12878 | 4170 | 4170 | 2985 | 1176 | 0.98 | 0.95 | 0.92 | 0.91 | 2795 | 805 |
| | HELA | 69 | 69 | 62 | 31 | 1 | 1 | 1 | 1 | 55 | 6 |
| | HMEC | 2558 | 2558 | 1907 | 735 | 0.99 | 0.98 | 0.97 | 0.96 | 1805 | 513 |
| | HUVEC | 163 | 163 | 143 | 54 | 1 | 1 | 1 | 1 | 137 | 19 |
| | IMR90 | 1009 | 1009 | 776 | 580 | 0.99 | 0.99 | 0.99 | 0.98 | 532 | 134 |
| | K562 | 1105 | 1105 | 859 | 344 | 1 | 1 | 0.99 | 0.99 | 796 | 211 |
| | NHEK | 5 | 5 | 5 | 1 | 1 | NA | 0 | NA | 5 | 0 |
| ChromHMM Gencode Rao cutoff 400 | GM12878 | 48128 | 14553 | 11493 | 5781 | 0.9 | 0.74 | 0.66 | 0.59 | 9625 | 2055 |
| ChromHMM Gencode Rao cutoff 300 | GM12878 | 69255 | 35135 | 25434 | 10692 | 0.87 | 0.73 | 0.62 | 0.58 | 23306 | 6390 |

| | | | | | | | | | | | |
|---|---|---|---|---|---|---|---|---|---|---|---|
| ChromHMM Gencode Rao cutoff 200 | GM12878 | 98562 | 63000 | 38936 | 14443 | 0.81 | 0.67 | 0.57 | 0.53 | 36885 | 13895 |
| ChromHMM Gencode Rao cutoff 150 | GM12878 | 122081 | 82931 | 48964 | 17056 | 0.8 | 0.67 | 0.56 | 0.53 | 46835 | 19051 |
| ChromHMM Gencode Rao cutoff 100 | GM12878 | 189085 | 150915 | 68712 | 22137 | 0.74 | 0.64 | 0.54 | 0.52 | 66953 | 36351 |
| | HMEC | 12773 | 1659 | 1541 | 1065 | 0.96 | 0.74 | 0.74 | 0.65 | 943 | 80 |
| | HUVEC | 2909 | 443 | 411 | 255 | 0.97 | 0.84 | 0.78 | 0.77 | 288 | 26 |
| | IMR90 | 12024 | 4995 | 4085 | 3032 | 0.95 | 0.85 | 0.87 | 0.8 | 2659 | 473 |
| | K562 | 47568 | 9723 | 8004 | 4295 | 0.91 | 0.73 | 0.66 | 0.6 | 6509 | 1224 |
| | NHEK | 46231 | 6062 | 5471 | 3435 | 0.95 | 0.76 | 0.74 | 0.66 | 3832 | 393 |
| ChromHMM Gencode Rao cutoff 50 | GM12878 | 465434 | 426121 | 106562 | 31765 | 0.64 | 0.6 | 0.49 | 0.48 | 105742 | 82607 |
| | HMEC | 46451 | 7232 | 6302 | 3911 | 0.94 | 0.77 | 0.74 | 0.65 | 4515 | 629 |
| | HUVEC | 39484 | 5793 | 5134 | 3032 | 0.95 | 0.77 | 0.72 | 0.67 | 3808 | 447 |
| | IMR90 | 24884 | 16998 | 12408 | 8588 | 0.9 | 0.76 | 0.79 | 0.71 | 9578 | 2574 |
| | K562 | 89227 | 42917 | 29077 | 11708 | 0.83 | 0.67 | 0.56 | 0.52 | 27161 | 8937 |
| | NHEK | 52543 | 11254 | 9528 | 5062 | 0.93 | 0.78 | 0.71 | 0.66 | 7671 | 1215 |
| Jin | IMR90 | 57578 | 50800 | 44239 | 8117 | 0.94 | 0.79 | 0.11 | 0.11 | 43107 | 5675 |
| FantomCage Gencode Jin | IMR90 | 1176 | 1167 | 743 | 401 | 0.9 | 0.84 | 0.77 | 0.73 | 641 | 260 |
| ChromHMM Gencode Jin | IMR90 | 5351 | 5303 | 3383 | 617 | 0.93 | 0.87 | 0.68 | 0.66 | 3323 | 1182 |
| FantomCage Gencode Chiapet | K562 | 2923 | 2916 | 1585 | 1869 | 0.8 | 0.75 | 0.86 | 0.75 | 902 | 491 |
| | MCF7 | 2190 | 2190 | 1471 | 1195 | 0.89 | 0.83 | 0.86 | 0.75 | 964 | 370 |
| ChromHMM Gencode Chiapet | K562 | 33598 | 33449 | 19550 | 6439 | 0.86 | 0.78 | 0.67 | 0.65 | 18868 | 8192 |
| FantomCage Gencode Javierre | Ery | 1562 | 74 | 44 | 64 | 1 | 1 | 1 | 1 | 14 | 6 |
| | Mac0 | 1352 | 88 | 59 | 64 | 0.98 | 0.94 | 0.98 | 0.95 | 29 | 12 |
| | Mac1 | 4077 | 215 | 144 | 153 | 1 | 1 | 1 | 1 | 66 | 36 |
| | Mac2 | 2305 | 112 | 75 | 85 | 0.99 | 0.96 | 0.98 | 0.96 | 33 | 10 |
| | MK | 1764 | 100 | 65 | 81 | 0.96 | 0.9 | 0.98 | 0.89 | 25 | 5 |
| | Mon | 2766 | 139 | 82 | 94 | 1 | 1 | 1 | 1 | 38 | 23 |
| | nCD4 | 1841 | 86 | 58 | 64 | 1 | 1 | 1 | 1 | 25 | 12 |
| | nCD8 | 1906 | 84 | 55 | 67 | 1 | 1 | 1 | 1 | 25 | 8 |
| | Neu | 3191 | 178 | 109 | 137 | 1 | 1 | 1 | 1 | 49 | 21 |
| ChromHMM Gencode Javierre | Ery | 77058 | 4484 | 2471 | 539 | 0.98 | 0.98 | 0.93 | 0.92 | 2408 | 1292 |
| | Mac0 | 34298 | 2003 | 1097 | 268 | 0.99 | 0.99 | 0.97 | 0.97 | 1074 | 568 |
| | Mac1 | 98273 | 4867 | 2996 | 658 | 0.97 | 0.96 | 0.91 | 0.9 | 2931 | 1300 |
| | Mac2 | 70264 | 3733 | 2298 | 474 | 0.99 | 0.99 | 0.95 | 0.94 | 2258 | 1021 |
| | MK | 66513 | 2629 | 1744 | 402 | 0.99 | 0.98 | 0.92 | 0.92 | 1691 | 630 |
| | Mon | 58426 | 2483 | 1547 | 330 | 0.96 | 0.94 | 0.91 | 0.9 | 1515 | 705 |
| | nCD4 | 51881 | 2975 | 1546 | 359 | 0.99 | 0.99 | 0.97 | 0.97 | 1510 | 838 |
| | nCD8 | 53476 | 2774 | 1623 | 339 | 0.98 | 0.97 | 0.93 | 0.93 | 1591 | 728 |
| | Neu | 97085 | 4661 | 2739 | 596 | 0.99 | 0.98 | 0.96 | 0.96 | 2678 | 1328 |

Table S2: BCC statistics for enhancers and promoters using randomly chosen enhancer-promoter pairs. Here the interactions between the enhancers and promoters are randomly assigned using the set of enhancers and promoters involved in the IEPs for the corresponding experimental condition. In this random assignment, the number of interactions for both enhancers and promoters were kept the same as in the original IEPs. Each of these random calculations was done 5 times and the average numbers are reported in this table. ChromHMM Gencode Rao cutoff 30 could not be listed due to longer processing time.

| Experiments | Cell lines | IEPs | IEPs >= 2.5kb | Enhancers | Promoters | BCC of enhancers | | BCC of promoters | | Enhancers with BCC>0 | Enhancers interacting with multiple promoters and BCC>0 |
|---|---|---|---|---|---|---|---|---|---|---|---|
| | | | | | | all | with multiple targets | all | with multiple targets | | |
| FantomCage Gencode Rao looplist | GM12878 | 434 | 434 | 317 | 286 | 0.62 | 0.17 | 0.38 | 0.29 | 3 | 1 |
| | HELA | 17 | 17 | 14 | 12 | NA | NA | NA | NA | 0 | 0 |
| | HMEC | 260 | 260 | 201 | 179 | 0.2 | 0.13 | 0.29 | 0.24 | 2 | 1 |
| | HUVEC | 19 | 19 | 17 | 13 | NA | NA | NA | NA | 0 | 0 |
| | IMR90 | 268 | 268 | 199 | 206 | 0.33 | 0.13 | 0.53 | 0.3 | 2 | 1 |
| | K562 | 89 | 89 | 70 | 70 | 0.2 | NA | 0.23 | 0.13 | 1 | 0 |
| | KBM7 | 8 | 8 | 5 | 8 | NA | NA | NA | NA | 0 | 0 |
| | NHEK | 1 | 1 | 1 | 1 | NA | NA | NA | NA | 0 | 0 |
| FantomCage Gencode Rao cutoff 400 | GM12878 | 3822 | 1231 | 997 | 949 | 0.68 | 0.39 | 0.63 | 0.35 | 25 | 8 |
| FantomCage Gencode Rao cutoff 300 | GM12878 | 5527 | 2861 | 2138 | 2092 | 0.56 | 0.32 | 0.56 | 0.34 | 135 | 56 |
| FantomCage Gencode Rao cutoff 200 | GM12878 | 8067 | 5240 | 3347 | 3328 | 0.45 | 0.29 | 0.47 | 0.3 | 437 | 236 |
| FantomCage Gencode Rao cutoff 150 | GM12878 | 10072 | 6938 | 4179 | 4199 | 0.41 | 0.27 | 0.45 | 0.28 | 731 | 434 |
| FantomCage Gencode Rao cutoff 100 | GM12878 | 16391 | 13223 | 6080 | 6788 | 0.28 | 0.21 | 0.38 | 0.24 | 2179 | 1596 |
| | HMEC | 734 | 124 | 113 | 116 | 0.3 | 0.1 | 0.5 | 0.1 | 1 | 1 |
| | HUVEC | 179 | 31 | 30 | 28 | NA | NA | NA | NA | 0 | 0 |
| | IMR90 | 2870 | 1056 | 885 | 877 | 0.72 | 0.43 | 0.72 | 0.42 | 23 | 7 |
| | K562 | 2677 | 618 | 524 | 503 | 0.56 | 0.42 | 0.68 | 0.41 | 6 | 3 |
| | KBM7 | 7152 | 1204 | 1112 | 1093 | 0.87 | 0.46 | 0.86 | 0.46 | 27 | 4 |
| | NHEK | 2901 | 425 | 387 | 385 | 0.8 | NA | 0.81 | 0.42 | 4 | 0 |
| FantomCage Gencode Rao cutoff 50 | GM12878 | 39558 | 36387 | 8906 | 13704 | 0.14 | 0.12 | 0.27 | 0.18 | 7107 | 6268 |
| | HMEC | 2896 | 511 | 453 | 456 | 0.63 | 0.45 | 0.76 | 0.42 | 5 | 2 |
| | HUVEC | 2422 | 454 | 401 | 392 | 0.65 | 0.2 | 0.77 | 0.46 | 5 | 1 |
| | IMR90 | 5589 | 3513 | 2455 | 2721 | 0.52 | 0.32 | 0.64 | 0.37 | 216 | 107 |
| | K562 | 5356 | 2820 | 2004 | 2101 | 0.51 | 0.32 | 0.6 | 0.35 | 135 | 65 |
| | KBM7 | 9191 | 2942 | 2489 | 2419 | 0.72 | 0.39 | 0.67 | 0.39 | 145 | 39 |
| | NHEK | 3348 | 780 | 672 | 659 | 0.67 | 0.45 | 0.76 | 0.39 | 15 | 5 |
| FantomCage Gencode Rao cutoff 30 | GM12878 | 70962 | 67791 | 9909 | 20636 | 0.08 | 0.08 | 0.2 | 0.15 | 9522 | 8956 |
| | HMEC | 4400 | 1718 | 1411 | 1380 | 0.67 | 0.41 | 0.66 | 0.38 | 47 | 16 |
| | HUVEC | 3910 | 1418 | 1148 | 1105 | 0.63 | 0.38 | 0.65 | 0.35 | 37 | 14 |
| | IMR90 | 10089 | 7951 | 4222 | 5348 | 0.33 | 0.25 | 0.53 | 0.32 | 912 | 637 |
| | K562 | 9138 | 6438 | 3458 | 4183 | 0.34 | 0.25 | 0.51 | 0.3 | 621 | 426 |
| | KBM7 | 13475 | 6608 | 5318 | 5043 | 0.66 | 0.38 | 0.61 | 0.35 | 708 | 236 |
| | NHEK | 5304 | 2448 | 1974 | 1946 | 0.66 | 0.38 | 0.66 | 0.39 | 97 | 35 |
| ChromHMM Gencode Rao looplist | GM12878 | 4170 | 4170 | 2985 | 1176 | 0.55 | 0.33 | 0.14 | 0.13 | 294 | 139 |
| | HELA | 69 | 69 | 62 | 31 | 0.2 | NA | NA | NA | 1 | 0 |
| | HMEC | 2558 | 2558 | 1907 | 735 | 0.58 | 0.37 | 0.13 | 0.13 | 105 | 49 |
| | HUVEC | 163 | 163 | 143 | 54 | 0.2 | 0.2 | NA | NA | 1 | 1 |
| | IMR90 | 1009 | 1009 | 776 | 580 | 0.57 | 0.4 | 0.38 | 0.29 | 23 | 9 |
| | K562 | 1105 | 1105 | 859 | 344 | 0.65 | 0.45 | 0.17 | 0.15 | 22 | 9 |
| | NHEK | 5 | 5 | 5 | 1 | NA | NA | NA | NA | 0 | 0 |
| ChromHMM Gencode Rao cutoff 400 | GM12878 | 48128 | 14553 | 11493 | 5781 | 0.66 | 0.38 | 0.21 | 0.17 | 3065 | 1004 |

| | | | | | | | | | | | |
|---|---|---|---|---|---|---|---|---|---|---|---|
| ChromHMM Gencode Rao cutoff 300 | GM12878 | 69255 | 35135 | 25434 | 10692 | 0.59 | 0.35 | 0.16 | 0.14 | 13618 | 5054 |
| ChromHMM Gencode Rao cutoff 200 | GM12878 | 98562 | 63000 | 38936 | 14443 | 0.48 | 0.3 | 0.12 | 0.11 | 29588 | 13127 |
| ChromHMM Gencode Rao cutoff 150 | GM12878 | 122081 | 82931 | 48964 | 17056 | 0.45 | 0.29 | 0.11 | 0.1 | 41238 | 18734 |
| ChromHMM Gencode Rao cutoff 100 | GM12878 | 189085 | 150915 | 68712 | 22137 | 0.33 | 0.23 | 0.08 | 0.07 | 66266 | 36700 |
| | HMEC | 12773 | 1659 | 1541 | 1065 | 0.89 | 0.47 | 0.38 | 0.26 | 48 | 6 |
| | HUVEC | 2909 | 443 | 411 | 255 | 0.52 | 0.2 | 0.08 | 0.08 | 3 | 1 |
| | IMR90 | 12024 | 4995 | 4085 | 3032 | 0.67 | 0.38 | 0.41 | 0.29 | 420 | 134 |
| | K562 | 47568 | 9723 | 8004 | 4295 | 0.69 | 0.4 | 0.25 | 0.2 | 1459 | 430 |
| | NHEK | 46231 | 6062 | 5471 | 3435 | 0.83 | 0.43 | 0.37 | 0.27 | 608 | 102 |
| ChromHMM Gencode Rao cutoff 50 | GM12878 | 465434 | 426121 | 106562 | 31765 | 0.16 | 0.13 | 0.04 | 0.04 | 106541 | 82905 |
| | HMEC | 46451 | 7232 | 6302 | 3911 | 0.78 | 0.42 | 0.34 | 0.25 | 845 | 183 |
| | HUVEC | 39484 | 5793 | 5134 | 3032 | 0.79 | 0.42 | 0.31 | 0.24 | 561 | 109 |
| | IMR90 | 24884 | 16998 | 12408 | 8588 | 0.58 | 0.35 | 0.32 | 0.24 | 3986 | 1604 |
| | K562 | 89227 | 42917 | 29077 | 11708 | 0.53 | 0.33 | 0.15 | 0.13 | 18005 | 7625 |
| | NHEK | 52543 | 11254 | 9528 | 5062 | 0.73 | 0.4 | 0.24 | 0.2 | 1942 | 490 |
| Jin | IMR90 | 57578 | 50800 | 44239 | 8117 | 0.81 | 0.44 | 0.09 | 0.08 | 44155 | 15977 |
| FantomCage Gencode Jin | IMR90 | 1176 | 1167 | 743 | 401 | 0.43 | 0.28 | 0.19 | 0.16 | 24 | 359 |
| ChromHMM Gencode Jin | IMR90 | 5351 | 5303 | 3383 | 617 | 0.45 | 0.29 | 0.05 | 0.05 | 446 | 1038 |
| FantomCage Gencode Chiapet | K562 | 2923 | 2916 | 1585 | 1869 | 0.33 | 0.26 | 0.47 | 0.3 | 140 | 1370 |
| | MCF7 | 2190 | 2190 | 1471 | 1195 | 0.5 | 0.32 | 0.33 | 0.22 | 76 | 1378 |
| ChromHMM Gencode Chiapet | K562 | 33598 | 33449 | 19550 | 6439 | 0.42 | 0.29 | 0.09 | 0.09 | 11156 | 8848 |
| FantomCage Gencode Javierre | Ery | 1562 | 74 | 44 | 64 | NA | NA | 0.4 | NA | 0 | 0 |
| | Mac0 | 1352 | 88 | 59 | 64 | 0.17 | 0.17 | 0.16 | 0.07 | 11 | 8 |
| | Mac1 | 4077 | 215 | 144 | 153 | 0.35 | 0.1 | 0.18 | 0.12 | 29 | 7 |
| | Mac2 | 2305 | 112 | 75 | 85 | NA | NA | 0.05 | 0.05 | 0 | 0 |
| | MK | 1764 | 100 | 65 | 81 | 0.17 | 0.17 | 0.35 | 0.1 | 1 | 157 |
| | Mon | 2766 | 139 | 82 | 94 | 0.11 | 0.11 | 0.27 | 0.07 | 16 | 16 |
| | nCD4 | 1841 | 86 | 58 | 64 | 0.2 | 0.2 | 0.2 | NA | 21 | 11 |
| | nCD8 | 1906 | 84 | 55 | 67 | 0.25 | 0.05 | 0.4 | NA | 286 | 143 |
| | Neu | 3191 | 178 | 109 | 137 | 0.07 | 0.07 | 0.14 | 0.14 | 1 | 1 |
| ChromHMM Gencode Javierre | Ery | 77058 | 4484 | 2471 | 539 | 0.34 | 0.28 | 0.05 | 0.03 | 8 | 6 |
| | Mac0 | 34298 | 2003 | 1097 | 268 | 0.38 | 0.28 | 0.03 | 0.03 | 9 | 7 |
| | Mac1 | 98273 | 4867 | 2996 | 658 | 0.42 | 0.31 | 0.05 | 0.05 | 10 | 7 |
| | Mac2 | 70264 | 3733 | 2298 | 474 | 0.42 | 0.31 | 0.04 | 0.04 | 7 | 5 |
| | MK | 66513 | 2629 | 1744 | 402 | 0.46 | 0.33 | 0.06 | 0.06 | 6 | 3 |
| | Mon | 58426 | 2483 | 1547 | 330 | 0.45 | 0.33 | 0.07 | 0.06 | 4 | 3 |
| | nCD4 | 51881 | 2975 | 1546 | 359 | 0.32 | 0.25 | 0.04 | 0.04 | 6 | 5 |
| | nCD8 | 53476 | 2774 | 1623 | 339 | 0.4 | 0.29 | 0.04 | 0.03 | 9 | 6 |
| | Neu | 97085 | 4661 | 2739 | 596 | 0.38 | 0.3 | 0.05 | 0.05 | 316 | 222 |

Table S3: Clusters of enhancers. Using the sharing enhancers (BCC > 0) we generated cluster of enhancers. An enhancer shares at least one promoter target with all the other enhancers in its cluster.

| | Cell lines | Enhancers | Enhancer clusters | Enhancers belong to the clusters | % of total enhancers belong to the clusters | Average number of enhancers in a cluster |
|---|---|---|---|---|---|---|
| FantomCage | GM12878 | 317 | 69 | 172 | 54.26 | 2.49 |
| | HELA | 14 | 2 | 6 | 42.86 | 3 |
| | HMEC | 201 | 39 | 101 | 50.25 | 2.59 |
| | HUVEC | 17 | 5 | 10 | 58.82 | 2 |
| | IMR90 | 199 | 36 | 82 | 41.21 | 2.28 |
| | K562 | 70 | 10 | 25 | 35.71 | 2.5 |
| | KBM7 | 5 | 0 | 0 | 0 | NA |
| | NHEK | 1 | 0 | 0 | 0 | NA |
| ChromHMM | GM12878 | 2985 | 653 | 2795 | 93.63 | 4.28 |
| | HELA | 62 | 20 | 54 | 87.1 | 2.7 |
| | HMEC | 1907 | 450 | 1805 | 94.65 | 4.01 |
| | HUVEC | 143 | 41 | 137 | 95.8 | 3.34 |
| | IMR90 | 776 | 200 | 531 | 68.43 | 2.65 |
| | K562 | 859 | 202 | 795 | 92.55 | 3.94 |
| | NHEK | 5 | 1 | 5 | 100 | 5 |
| FantomCage (cutoffs) | GM12878 | 997 | 159 | 382 | 38.31 | 2.4 |
| | HMEC | 113 | 8 | 16 | 14.16 | 2 |
| | HUVEC | 30 | 3 | 6 | 20 | 2 |
| | IMR90 | 885 | 130 | 280 | 31.64 | 2.15 |
| | K562 | 524 | 72 | 168 | 32.06 | 2.33 |
| | KBM7 | 1112 | 96 | 203 | 18.26 | 2.11 |
| | NHEK | 387 | 35 | 75 | 19.38 | 2.14 |
| ChromHMM (cutoffs) | GM12878 | 11493 | 2755 | 9624 | 83.74 | 3.49 |
| | HMEC | 1541 | 386 | 943 | 61.19 | 2.44 |
| | HUVEC | 411 | 112 | 287 | 69.83 | 2.56 |
| | IMR90 | 4085 | 1031 | 2658 | 65.07 | 2.58 |
| | K562 | 8004 | 2076 | 6509 | 81.32 | 3.14 |
| | NHEK | 5471 | 1433 | 3832 | 70.04 | 2.67 |

Here we used the IEPs with FantomCage and ChromHMM enhancers and Gencode promoters using Rao looplists and cutoffs (400 for GM12878 and 100 for other cell lines.)

Table S4: The distance between each consecutive enhancer pairs in an enhancer clusters are shown in the left columns of the table. The right columns show the distance between each consecutive target pairs of the targets of the enhancers in an enhancer cluster.

| | Cell lines | Enhancer clusters | **Distance distribution between the enhancers of the same clusters** | | | | | | **Distance distribution between the targets of enhancers of the same clusters** | | | | | |
|---|---|---|---|---|---|---|---|---|---|---|---|---|---|---|
| | | | <=1kb | > 1kb and <= 5kb | > 5kb and <= 10kb | > 10kb and <= 50kb | > 50kb | Diff Chrom | <=1kb | > 1kb and <= 5kb | > 5kb and <= 10kb | > 10kb and <= 50kb | > 50kb | Diff Chrom |
| FantomCage | GM12878 | 69 | 48.72 | 40.37 | 0 | 2.9 | 8.02 | 0 | 61.54 | 11.54 | 0 | 0 | 26.92 | 0 |
| | HELA | 2 | 58.34 | 41.66 | 0 | 0 | 0 | 0 | 100 | 0 | 0 | 0 | 0 | 0 |
| | HMEC | 39 | 49.57 | 34.02 | 0 | 5.81 | 10.6 | 0 | 64.29 | 14.29 | 0 | 0 | 21.43 | 0 |
| | HUVEC | 5 | 60 | 40 | 0 | 0 | 0 | 0 | 100 | 0 | 0 | 0 | 0 | 0 |
| | IMR90 | 36 | 70.83 | 20.83 | 0 | 0 | 8.33 | 0 | 77.78 | 14.81 | 0 | 0 | 7.41 | 0 |
| | K562 | 10 | 65 | 25 | 0 | 10 | 0 | 0 | 66.67 | 33.33 | 0 | 0 | 0 | 0 |
| | KBM7 | 0 | NA | NA | NA | NA | NA | NA | NA | NA | NA | NA | NA | NA |
| | NHEK | 0 | NA | NA | NA | NA | NA | NA | NA | NA | NA | NA | NA | NA |
| ChromHMM | GM12878 | 653 | 63.69 | 28.67 | 0 | 0.83 | 6.8 | 0 | 55.59 | 18.81 | 0.06 | 1.52 | 24.02 | 0 |
| | HELA | 20 | 65.17 | 34.83 | 0 | 0 | 0 | 0 | 50 | 50 | 0 | 0 | 0 | 0 |
| | HMEC | 450 | 67.21 | 28.88 | 0 | 0.81 | 3.1 | 0 | 70.57 | 17.76 | 0 | 1.02 | 10.65 | 0 |
| | HUVEC | 41 | 77.97 | 22.03 | 0 | 0 | 0 | 0 | 66.67 | 33.33 | 0 | 0 | 0 | 0 |
| | IMR90 | 200 | 73.16 | 19.49 | 0 | 0 | 7.35 | 0 | 68.63 | 22.88 | 0 | 0 | 8.5 | 0 |
| | K562 | 202 | 70.74 | 27.03 | 0 | 0.14 | 2.09 | 0 | 61.9 | 31.55 | 0 | 1.79 | 4.76 | 0 |
| | NHEK | 1 | 50 | 50 | 0 | 0 | 0 | 0 | NA | NA | NA | NA | NA | NA |
| FantomCage (cutoffs) | GM12878 | 159 | 49.87 | 32.87 | 9.04 | 8.22 | 0 | 0 | 48.72 | 22.91 | 12.13 | 16.24 | 0 | 0 |
| | HMEC | 8 | 75 | 0 | 25 | 0 | 0 | 0 | 33.33 | 0 | 66.67 | 0 | 0 | 0 |
| | HUVEC | 3 | 100 | 0 | 0 | 0 | 0 | 0 | 47.92 | 24.17 | 13.33 | 14.58 | 0 | 0 |
| | IMR90 | 130 | 62.26 | 18 | 9.23 | 10.51 | 0 | 0 | 64.29 | 15.87 | 14.29 | 5.56 | 0 | 0 |
| | K562 | 72 | 60.14 | 27.22 | 10.79 | 1.85 | 0 | 0 | 89.74 | 0 | 5.13 | 5.13 | 0 | 0 |
| | KBM7 | 96 | 57.99 | 10.07 | 29.86 | 1.04 | 1.04 | 0 | 100 | 0 | 0 | 0 | 0 | 0 |
| | NHEK | 35 | 80 | 0 | 20 | 0 | 0 | 0 | 65.52 | 29.45 | 5.02 | 0 | 0 | 0 |
| ChromHMM (cutoffs) | GM12878 | 2755 | 70.14 | 14.85 | 9.43 | 5.53 | 0.05 | 0 | 41.27 | 16.24 | 21.06 | 21.06 | 0.36 | 0 |
| | HMEC | 386 | 91.78 | 1.37 | 6.7 | 0.16 | 0 | 0 | 77.78 | 0 | 22.22 | 0 | 0 | 0 |
| | HUVEC | 112 | 91.77 | 5.19 | 2.59 | 0.45 | 0 | 0 | 76.92 | 0 | 0 | 0 | 23.08 | 0 |
| | IMR90 | 1031 | 80.53 | 7.31 | 5.5 | 6.57 | 0.1 | 0 | 43.36 | 19.4 | 15.55 | 21.32 | 0.37 | 0 |
| | K562 | 2076 | 75.94 | 11.21 | 8.85 | 3.83 | 0.17 | 0 | 43.01 | 14.27 | 23.42 | 17.82 | 1.47 | 0 |
| | NHEK | 1433 | 88.4 | 2.24 | 9.24 | 0.03 | 0.08 | 0 | 71.77 | 0.19 | 27.04 | 0 | 1 | 0 |

Here we used the IEPs with FantomCage and ChromHMM enhancers and Gencode promoters using Rao looplists and cutoffs (400 for GM12878 and 100 for other cell lines.)

Table S5: Overlap between enhancer clusters and super-enhancers. The percentage of enhancer clusters overlapped with the super-enhancers and the percentage of super-enhancers overlapped with the enhancer clusters are shown. The right columns of the table show that, among the percentage of enhancer clusters overlapping with the super-enhancers (5th column), what percentage of enhancer clusters are involved in different amount of region overlap. This shows on average whether a cluster have a high or low region overlap with a super-enhancer.

| | Cell lines | Number of clusters | Number of super-enhancers | % of clusters overlapped with super-enhancers | % of super enhancers overlapped with clusters | Distribution of the percentage of overlap between enhancer clusters and super-enhancers | | | |
|---|---|---|---|---|---|---|---|---|---|
| | | | | | | <= 25% | > 25% and <= 50% | > 50% and <= 75% | > 75% and <= 100% |
| FantomCage | GM12878 | 69 | 257 | 27.54 | 7.39 | 4.35 | 4.35 | 1.45 | 17.39 |
| | HELA | 2 | 698 | 100 | 0.29 | 0 | 0 | 0 | 100 |
| | HMEC | 39 | 1099 | 51.28 | 2.46 | 12.82 | 2.56 | 2.56 | 33.33 |
| | HUVEC | 5 | 912 | 40 | 0.22 | 0 | 0 | 0 | 40 |
| | IMR90 | 36 | 502 | 30.56 | 2.79 | 5.56 | 0 | 2.78 | 22.22 |
| | K562 | 10 | 742 | 50 | 0.67 | 0 | 0 | 10 | 40 |
| | NHEK | 0 | 1024 | NA | NA | NA | NA | NA | NA |
| ChromHMM | GM12878 | 653 | 257 | 9.8 | 23.35 | 1.68 | 1.38 | 0.61 | 6.13 |
| | HELA | 20 | 698 | 30 | 0.72 | 0 | 0 | 0 | 30 |
| | HMEC | 450 | 1099 | 21.56 | 9.37 | 2.67 | 0.89 | 1.56 | 16.44 |
| | HUVEC | 41 | 912 | 34.15 | 1.54 | 0 | 0 | 0 | 34.15 |
| | IMR90 | 200 | 502 | 12.5 | 6.77 | 4.5 | 1 | 0 | 7 |
| | K562 | 202 | 742 | 17.82 | 4.99 | 1.49 | 0.99 | 2.97 | 12.38 |
| | NHEK | 1 | 1024 | 0 | 0 | 0 | 0 | 0 | 0 |
| FantomCage (cutoffs) | GM12878 | 159 | 257 | 20.75 | 12.06 | 0 | 0 | 0 | 20.75 |
| | HMEC | 8 | 1099 | 25 | 0.18 | 0 | 0 | 0 | 25 |
| | HUVEC | 3 | 912 | 33.33 | 0.11 | 0 | 0 | 0 | 33.33 |
| | IMR90 | 130 | 502 | 19.23 | 4.78 | 0 | 0 | 0.77 | 18.46 |
| | K562 | 72 | 742 | 36.11 | 3.1 | 1.39 | 0 | 1.39 | 33.33 |
| | NHEK | 35 | 1024 | 34.29 | 1.17 | 0 | 0 | 0 | 34.29 |
| ChromHMM (cutoffs) | GM12878 | 2755 | 257 | 7.7 | 45.91 | 0.4 | 0.18 | 0.62 | 6.5 |
| | HMEC | 386 | 1099 | 16.84 | 5.46 | 1.55 | 0.26 | 1.04 | 13.99 |
| | HUVEC | 112 | 912 | 16.07 | 1.97 | 0 | 0.89 | 0 | 15.18 |
| | IMR90 | 1031 | 502 | 5.82 | 9.96 | 0.29 | 0.29 | 0.1 | 5.14 |
| | K562 | 2076 | 742 | 13.68 | 29.51 | 1.25 | 1.78 | 1.01 | 9.63 |
| | NHEK | 1433 | 1024 | 13.47 | 17.29 | 0.42 | 0.49 | 1.4 | 11.17 |

Here we used the IEPs with FantomCage and ChromHMM enhancers and Gencode promoters using Rao looplists and cutoffs (400 for GM12878 and 100 for other cell lines.)

Table S6: Average percentage of enhancers in the same clusters mapped in a common TAD or TAD gap. We mapped the enhancer clusters to the defined TADs and the gaps between two TADs. On average, almost all of the enhancers in a cluster were found to be located within the same TAD or TAD gap.

| | Cell lines | Enhancer clusters | Relevant enhancer clusters (having at least 2 enhancers assigned to tads) | % of relevant enhancer clusters belonging to common tad | Relevant enhancer clusters (having at least 2 enhancers assigned to tad gap) | % of relevant enhancer clusters belonging to common tad gap |
|---|---|---|---|---|---|---|
| FantomCage | GM12878 | 69 | 66 | 98.48 | 14 | 100 |
| | HELA | 2 | 0 | 0 | 2 | 100 |
| | HMEC | 39 | 24 | 100 | 17 | 94.12 |
| | HUVEC | 5 | 3 | 100 | 2 | 100 |
| | IMR90 | 36 | 35 | 100 | 5 | 100 |
| | K562 | 10 | 8 | 100 | 1 | 100 |
| | KBM7 | 0 | 0 | 0 | 0 | 0 |
| | NHEK | 0 | 0 | 0 | 0 | 0 |
| | IMR90 (Dixon) | 36 | 32 | 100 | 4 | 100 |
| ChromHMM | GM12878 | 653 | 619 | 98.38 | 170 | 97.65 |
| | HELA | 20 | 15 | 100 | 6 | 100 |
| | HMEC | 450 | 295 | 100 | 180 | 98.89 |
| | HUVEC | 41 | 33 | 100 | 12 | 100 |
| | IMR90 | 200 | 184 | 99.46 | 36 | 100 |
| | K562 | 202 | 159 | 100 | 40 | 100 |
| | NHEK | 1 | 0 | 0 | 1 | 100 |
| | IMR90 (Dixon) | 200 | 193 | 99.48 | 8 | 100 |
| FantomCage (cutoffs) | GM12878 | 159 | 143 | 100 | 27 | 100 |
| | HMEC | 8 | 5 | 100 | 3 | 100 |
| | HUVEC | 3 | 2 | 100 | 1 | 100 |
| | IMR90 | 130 | 110 | 100 | 26 | 100 |
| | K562 | 72 | 59 | 100 | 16 | 100 |
| | KBM7 | 96 | 79 | 100 | 21 | 100 |
| | NHEK | 35 | 20 | 100 | 15 | 100 |
| | IMR90 (Dixon) | 130 | 120 | 100 | 10 | 100 |
| ChromHMM (cutoffs) | GM12878 | 2755 | 2347 | 99.79 | 758 | 100 |
| | HMEC | 386 | 194 | 100 | 199 | 100 |
| | HUVEC | 112 | 58 | 100 | 57 | 100 |
| | IMR90 | 1031 | 866 | 99.88 | 208 | 100 |
| | K562 | 2076 | 1573 | 99.87 | 651 | 100 |
| | NHEK | 1433 | 890 | 100 | 592 | 100 |
| | IMR90 (Dixon) | 1031 | 936 | 100 | 98 | 100 |

Here we used the IEPs with FantomCage and ChromHMM enhancers and Gencode promoters using Rao looplists and cutoffs (400 for GM12878 and 100 for other cell lines.)

Table S7: Percentages of common enhancer clusters between two cell lines.

| | Cell line1 | Cell line2 | Clusters in cell line 1 | Clusters in cell line 2 | % of common clusters (with respect to cell line 1) | % of common clusters (with respect to cell line 2) |
|---|---|---|---|---|---|---|
| FantomCage | GM12878 | HELA | 69 | 2 | 0 | 0 |
| | GM12878 | HMEC | 69 | 39 | 1.45 | 2.56 |
| | GM12878 | HUVEC | 69 | 5 | 0 | 0 |
| | GM12878 | IMR90 | 69 | 36 | 2.9 | 5.56 |
| | GM12878 | K562 | 69 | 10 | 5.8 | 40 |
| | GM12878 | KBM7 | 69 | 0 | NA | NA |
| | GM12878 | NHEK | 69 | 0 | NA | NA |
| | HELA | HMEC | 2 | 39 | 50 | 2.56 |
| | HELA | HUVEC | 2 | 5 | 0 | 0 |
| | HELA | IMR90 | 2 | 36 | 0 | 0 |
| | HELA | K562 | 2 | 10 | 0 | 0 |
| | HELA | KBM7 | 2 | 0 | NA | NA |
| | HELA | NHEK | 2 | 0 | NA | NA |
| | HMEC | HUVEC | 39 | 5 | 0 | 0 |
| | HMEC | IMR90 | 39 | 36 | 10.26 | 11.11 |
| | HMEC | K562 | 39 | 10 | 2.56 | 10 |
| | HMEC | KBM7 | 39 | 0 | NA | NA |
| | HMEC | NHEK | 39 | 0 | NA | NA |
| | HUVEC | IMR90 | 5 | 36 | 20 | 2.78 |
| | HUVEC | K562 | 5 | 10 | 20 | 10 |
| | HUVEC | KBM7 | 5 | 0 | NA | NA |
| | HUVEC | NHEK | 5 | 0 | NA | NA |
| | IMR90 | K562 | 36 | 10 | 5.56 | 20 |
| | IMR90 | KBM7 | 36 | 0 | NA | NA |
| | IMR90 | NHEK | 36 | 0 | NA | NA |
| | K562 | KBM7 | 10 | 0 | NA | NA |
| | K562 | NHEK | 10 | 0 | NA | NA |
| | KBM7 | NHEK | 0 | 0 | NA | NA |
| ChromHMM | GM12878 | HELA | 653 | 20 | 0 | 0 |
| | GM12878 | HMEC | 653 | 450 | 1.23 | 1.78 |
| | GM12878 | HUVEC | 653 | 41 | 0.15 | 2.44 |
| | GM12878 | IMR90 | 653 | 200 | 0.46 | 1.5 |
| | GM12878 | K562 | 653 | 202 | 1.23 | 3.96 |
| | GM12878 | KBM7 | 653 | 1 | 0 | 0 |
| | HELA | HMEC | 20 | 450 | 5 | 0.22 |
| | HELA | HUVEC | 20 | 41 | 0 | 0 |
| | HELA | IMR90 | 20 | 200 | 0 | 0 |
| | HELA | K562 | 20 | 202 | 15 | 1.49 |
| | HELA | KBM7 | 20 | 1 | 0 | 0 |
| | HMEC | HUVEC | 450 | 41 | 0.44 | 4.88 |
| | HMEC | IMR90 | 450 | 200 | 1.33 | 3 |
| | HMEC | K562 | 450 | 202 | 1.11 | 2.48 |
| | HMEC | KBM7 | 450 | 1 | 0 | 0 |
| | HUVEC | IMR90 | 41 | 200 | 4.88 | 1 |
| | HUVEC | K562 | 41 | 202 | 2.44 | 0.5 |
| | HUVEC | KBM7 | 41 | 1 | 0 | 0 |
| | IMR90 | K562 | 200 | 202 | 1 | 0.99 |
| | IMR90 | KBM7 | 200 | 1 | 0 | 0 |
| | K562 | KBM7 | 202 | 1 | 0 | 0 |
| FantomCage (cutoffs) | GM12878 | HELA | 159 | 8 | 3.14 | 62.5 |
| | GM12878 | HMEC | 159 | 3 | 0.63 | 33.33 |
| | GM12878 | HUVEC | 159 | 130 | 19.5 | 23.85 |
| | GM12878 | IMR90 | 159 | 72 | 27.04 | 59.72 |
| | GM12878 | K562 | 159 | 96 | 22.01 | 36.46 |
| | GM12878 | KBM7 | 159 | 35 | 12.58 | 57.14 |
| | HELA | HMEC | 8 | 3 | 12.5 | 33.33 |
| | HELA | HUVEC | 8 | 130 | 62.5 | 3.85 |
| | HELA | IMR90 | 8 | 72 | 75 | 8.33 |
| | HELA | K562 | 8 | 96 | 100 | 8.33 |
| | HELA | KBM7 | 8 | 35 | 75 | 17.14 |
| | HMEC | HUVEC | 3 | 130 | 66.67 | 1.54 |
| | HMEC | IMR90 | 3 | 72 | 33.33 | 1.39 |

| | | | | | | |
|---|---|---|---|---|---|---|
| | HMEC | K562 | 3 | 96 | 100 | 3.12 |
| | HMEC | KBM7 | 3 | 35 | 33.33 | 2.86 |
| | HUVEC | IMR90 | 130 | 72 | 18.46 | 33.33 |
| | HUVEC | K562 | 130 | 96 | 15.38 | 20.83 |
| | HUVEC | KBM7 | 130 | 35 | 10.77 | 40 |
| | IMR90 | K562 | 72 | 96 | 41.67 | 31.25 |
| | IMR90 | KBM7 | 72 | 35 | 25 | 51.43 |
| | K562 | KBM7 | 96 | 35 | 36.46 | 100 |
| ChromHMM (cutoffs) | GM12878 | HELA | 2755 | 386 | 0.58 | 4.15 |
| | GM12878 | HMEC | 2755 | 112 | 0.18 | 4.46 |
| | GM12878 | HUVEC | 2755 | 1031 | 0.83 | 2.23 |
| | GM12878 | IMR90 | 2755 | 2076 | 4.07 | 5.11 |
| | GM12878 | K562 | 2755 | 1433 | 1.71 | 3.21 |
| | HELA | HMEC | 386 | 112 | 1.81 | 6.25 |
| | HELA | HUVEC | 386 | 1031 | 2.85 | 1.07 |
| | HELA | IMR90 | 386 | 2076 | 6.99 | 1.35 |
| | HELA | K562 | 386 | 1433 | 6.48 | 1.74 |
| | HMEC | HUVEC | 112 | 1031 | 6.25 | 0.68 |
| | HMEC | IMR90 | 112 | 2076 | 9.82 | 0.63 |
| | HMEC | K562 | 112 | 1433 | 5.36 | 0.42 |
| | HUVEC | IMR90 | 1031 | 2076 | 3.1 | 1.59 |
| | HUVEC | K562 | 1031 | 1433 | 3.2 | 2.23 |
| | IMR90 | K562 | 2076 | 1433 | 2.75 | 3.98 |

Here we used the IEPs with FantomCage and ChromHMM enhancers and Gencode promoters using Rao looplists and cutoffs (400 for GM12878 and 100 for other cell lines.)
